# Single Molecule Mass Photometry Reveals Dynamic Oligomerization of Plant and Human Peroxiredoxins for Functional Conservation and Diversification

**DOI:** 10.1101/2021.01.30.428949

**Authors:** Michael Liebthal, Manish Singh Kushwah, Philipp Kukura, Karl-Josef Dietz

**Affiliations:** Department of Biochemistry and Physiology of Plants, Faculty of Biology, University of Bielefeld, 33615 Bielefeld, Germany; Physical and Theoretical Chemistry Laboratory, Department of Chemistry, University of Oxford, South Parks Road, Oxford OX1 3QZ, UK

**Keywords:** Arabidopsis thaliana, Homo sapiens, hydrogen peroxide, oligomerization, peroxiredoxin, redox state, single molecule counting

## Abstract

Single molecule mass photometry was used to study the dynamic equilibria of the ubiquitous and highly abundant 2-Cysteine peroxiredoxins (2-CysPRX). 2-CysPRXs adopt distinct functions in all cells dependent on their oligomeric conformation ranging from dimers to decamers and high molecular weight aggregates (HMW). The oligomeric state depends on the redox state of their catalytic cysteinyl residues. To which degree they interconvert, how the interconversion is regulated, and how the oligomerisation propensity is organism specific remains, however, poorly understood. The dynamics differs between wild-type and single point mutants affecting the oligomerization interfaces, with concomitant changes to function. Titrating concentration and redox state of *Arabidopsis thaliana* and human 2-CysPRXs revealed features conserved among all 2-CysPRX and clear differences concerning oligomer transitions, the occurrence of transition states and the formation of HMW which are associated with chaperone activity or storage. The results indicate functional differentiation of human 2-CysPRXs. Our results point to a diversified functionality of oligomerization for 2-CysPRXs and illustrate the power of mass photometry to non-invasively quantify oligomer distributions in a redox environment. This knowledge is important to fully address and model PRX function in cell redox signaling e.g., in photosynthesis, cardiovascular and neurological diseases or carcinogenesis.

## 1. Introduction

2-Cysteine peroxiredoxins (2-CysPRX) are the largest subgroup in the family of peroxiredoxins (PRX) and exhibit different subcellular and cellular localization. They fulfill diverse functions across the kingdoms of life and within the same cell depending on their redox state and subcellular localization. [1, 2, 3] Initially mainly considered as thiol peroxidases to scavenge reactive oxygen species (ROS), their accepted redox-dependent functions range from cell differentiation, tumor suppression, signal transduction, thioredoxin (TRX) oxidation and chaperone-dependent protein homeostasis to participation in stress resistance and disease control. [4, 5, 6, 7, 8]

The polydispersity of these enzymes, which is fundamentally linked to their thiol redox state, is associated with changes in their function – whether they are acting as a chaperone (hyperoxidized decamer, including high molecular weight aggregates), a polydisperse peroxidase (mixture of reduced dimers and decamers) or a redox signaling element (oxidized dimer) **(Figure 1A)**. [9] Other types of PRX either lack oligomeric conformations (PRXQ/bacterioferritin comigratory protein BCP) or adopt different quaternary structures (PRX type II). [2] Oligomer formation and dissociation thus control protein activity, protein turnover and exposure of interaction surfaces. [10] This complexity is enhanced even further in fungi, mammals and humans, where more than one 2-CysPRX are known, e.g. in humans where four 2-CysPRXs have been identified. [11, 12] In addition, they localize to different subcellular compartments like cytosol (HsPRX1, HsPRX2) and mitochondrion (HsPRX3) indicating distinct functions in metabolism and regulation.

**Figure 1:**
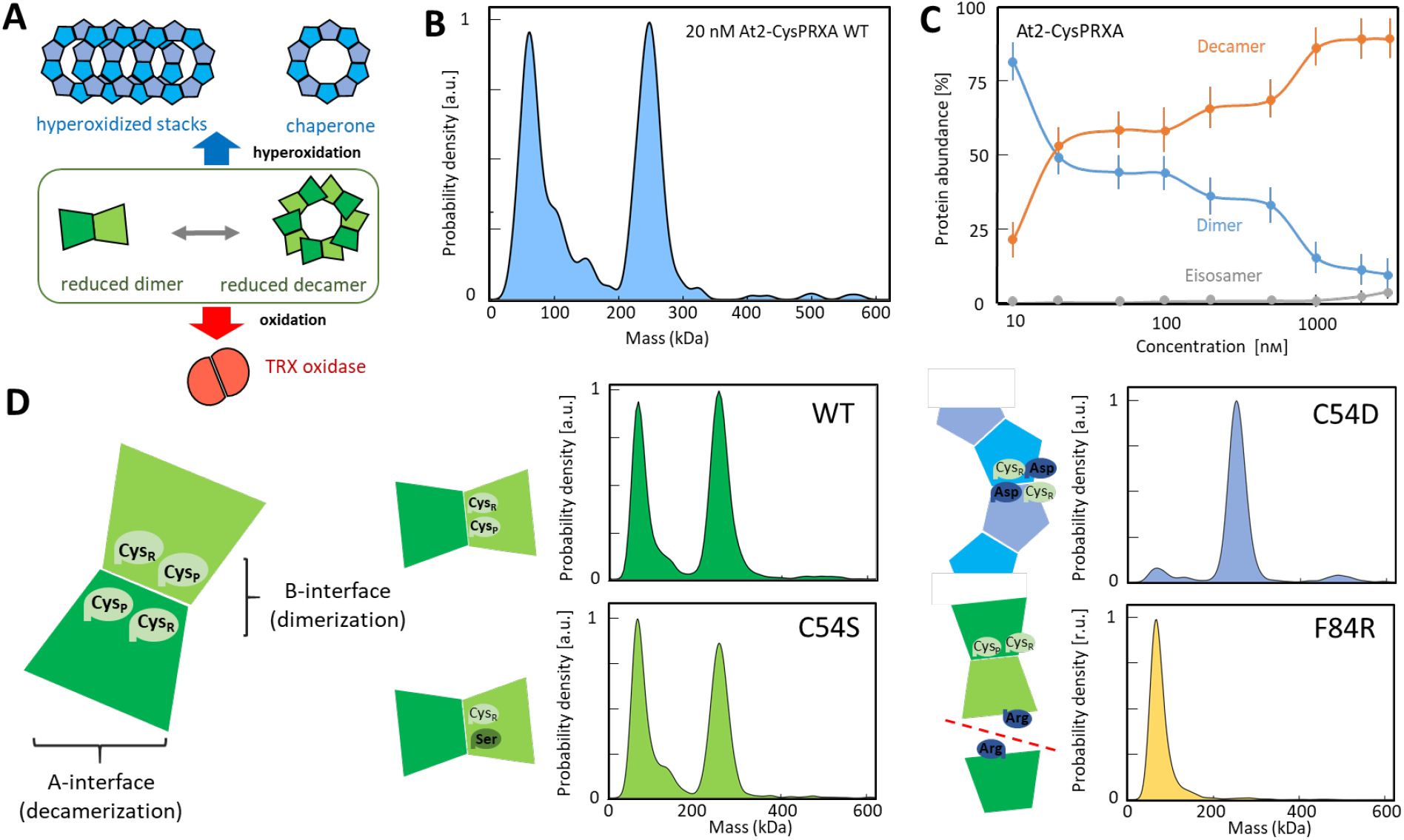
Schematics of the redox-dependent conformations adopted by the peroxiredoxin At2-CysPRXA and oligomer distribution analysis by single molecule mass photometry in dependence on concentration and after site-directed mutation. (A) At2-CysPRXA in dependence on its redox state adopts the dimeric, roughly globular conformation of 48 kDa (oxidized form [red], fraction of reduced form [green]), the doughnut-like decameric conformation of 240 kDa (fraction of reduced form [green ring], hyperoxidized form [blue ring]) or the form of hyperaggregates of *n*·240 kDa (hyperoxidized stacks of blue rings, with n being the number of stacked decamers). (B) Oligomerization state of At2-CysPRXA at 20 nm concentration as determined by mass photometry. Dimers (48 kDa) and decamers (240 kDa) were equally represented at this low concentration. (C) Oligomerization state of At2-CysPRXA as a function of concentration between 10 nm and 3 μm. (D) Conformational state of site-directed mutated variants of 2-CysPrxA at 100 nm concentration. The schematics illustrates the kind of mutation introduced into the protein. The monomers display two interfaces for protein-protein-interactions. The monomer/monomer-interface (B-interface) facilitates the dimerization and the dimer/dimer interface (A-interface) enables decamerization. Each monomer has a peroxidatic (CysP) and a resolving cysteine (CysR). Replacement of CysP by Ser (C54D) mimics the constitutively reduced form. Introducing the charged Asp in place of CysP mimics hyperoxidation of the Cys thiol to sulfinic acid (C54D: pseudo-hyperoxidized). Substitution of Arg for Phe (F84R) in the A-interface impedes decamerization. Mass photometry revealed that the pseudo-hyperoxidized C54D adopted the decameric form. Dimers and higher order aggregates were present at very low amounts. Statistics: All experiments were performed with *n>6* determinations on at least two different days. Reproducibility of all readings is shown in Supplemental Figure 2.

Its vast range of oligomeric states, ranging from dimers to large aggregates, makes PRX a particularly challenging target for functional studies. The different oligomeric states of 2-CysPRX have been observed and characterized by a variety of bioanalytical techniques. [13, 14, 15, 16, 17, 18] Each of these approaches has drawbacks, like requirements for high concentrations (size exclusion chromatography: SEC, dynamic light scattering: DLS), bulky labeling (fluorescence resonance energy transfer: FRET), poorly defined or non-native conditions (electron microscopy: EM, polyacrylamide gel electrophoresis: PAGE), or complexity of data interpretation (isothermal titration microcalorimetry: ITC). Taken together, these studies have shown beyond doubt that PRX is polydisperse and that changes in its oligomeric distributions can be traced to its function.

At the same time, our understanding of the molecular details of PRX polydispersity beyond fairly general observa tions, and its dependence on redox conditions, changes to the oligomerization interfaces and ultimately function, remains limited. The challenging questions concern (i) the precise distribution profile of the dimeric and oligomeric states and its dependence on structural features, (ii) the dynamics of transitions between the states in a constant physicochemical environment, (iii) the kinetics of conformational transitions depending on signaling cues like changing redox conditions and ROS and (iv) the effect of interactors. An example of open questions may be given for (ii) dynamics of transition: Do intermediates of multiple dimers occur at detectable concentrations before the decamer is formed, or are the dimer and decamer the only thermodynamically preferred long-lived states? These parameters determine 2-CysPRX function in the aforementioned cellular processes. To address these open questions, we applied single molecule mass photometry to reveal the oligomeric behavior of 2-CysPRXs for static and dynamic physicochemical environments and selected single point mutants from different organisms. [19]

## 2. Results

### 2.1. Concentration-dependent oligomerization of 2-CysPRX

Mass photometry essentially counts single molecules of defined masses appearing or leaving a detection plane, most commonly defined by a microscope cover glass surface. The first set of experiments tested the suitability of mass photometry to scrutinize the oligomerization state of 2-CysPRX. A low concentration of reduced Arabidopsis 2-CysPRXA (20 nm) revealed dimers (48 kDa) and decamers (240 kDa) detected as dominant species of equal intensity. Importantly, intermediate oligomers (tetramer, hexamer and so on) appeared as distinct peaks as well but their object counts decreased exponentially with size **(Figure 1B)**. This means that about five times more dimer was associated with the decameric fraction than with the dimer fraction under these conditions (**Supporting Figure 1**). Detected HMW aggregates had very low abundance. The distribution was highly reproducible and illustrates the purity of the protein solution (**Supporting Figure 2**). The detection limit of mass photometry is near 40 kDa and implies that we could not detect monomeric species, which is unlikely to be limiting in this case, given that dimers have consistently been reported as the minimal building block of 2-CysPRXs in the literature (reviewed in [2]).

Monitoring the oligomerization propensity with increasing concentrations ranging from 20 nm to 3 μm for At2-CysPRXA revealed unique oligomer dynamics leading to increased association of decamers and larger oligomers **(Figure 1C)**. Distinct oligomer populations were detected at around 500 kDa pointing towards stacked decamers or eicosamers even under reducing conditions. Their abundance increased at higher concentrations. The associated K_d_ for single components could not be determined due to the need for operation closer to physiological concentrations (30-60 μm, depending on organism), limitations of mass photometry at concentrations above 3 μm and unexpected oligomerization dynamics of At2-CysPRXA.

### 2.2. Site-directed mutations force 2-CysPRX into a specific conformation

The next experiments scrutinized the possibility to manipulate oligomerization by site-directed mutagenesis. Amino acid substitutions were intended to mimic specific conformations and to address the redox dependence of the oligomer/dimer-equilibria **(Figure 1D)**. [20] Specifically, we stabilized the reduced conformation by mutating the peroxidatic C54 to serine preventing disulfide formation. The C54S variant thus exhibited a small decrease in decamer abundance.

Substituting Asp for C54 mimics the hyperoxidized and charged sulfinyl group and stabilized decamers and hyperaggregates with chaperone function. Conversely, introducing the charged arginyl residue at the dimer-dimer interface in the F84R variant inhibited decamer formation completely, causing almost exclusive accumulation of dimers and minimal abundance of tetramers. Dynamic light scattering confirmed the mass photometry results at higher protein concentrations of 10 μm, albeit at much lower mass resolution (**Supporting Figure 3**). The oligomerization state thus sensitively responded to the intentionally introduced changes in amino acid side chains suggesting a delicate interplay between oligomerization and structural features. The precision of the single mass photometry result surpassed the rather vague result from SEC. [21]

### 2.3. Thiol-modifications control oligomerization state

The redox state of 2-CysPRX strongly affects oligomerization through the peroxidatic cysteine (CysP54) reacting with H_2_O_2_ and forming a disulfide bond with the resolving cysteine (CysR176). CysP together with vicinal amino acid residues Arg, Thr, and Pro forms a catalytic pocket in the fully folded (FF) structure. [22] This molecular environment lowers the pKa-value of CysP below 6 under physiological conditions giving rise to an extraordinarily high reactivity towards peroxide substrates. [23] Upon oxidation, the protein conformation changes to the locally unfolded (LU) state allowing for disulfide formation with CysR, which exhibits a weakened dimer-dimer interface causing oligomer dissociation. [1]

In line with this model, we found that exposing At2-CysPRXA to H_2_O_2_ caused a significant shift away from decamers toward dimers **(Figure 2A)**. By contrast, simultaneous exposure to both H_2_O_2_ and dithiothreitol (DTT), which promotes repeated catalytic turnover and stimulates hyperoxidation of the C54 thiol to a sulfinic acid residue, led to an increase in the decamer-to-dimer ratio beyond that observed in the reduced state (Figure 2A).

**Figure 2.**
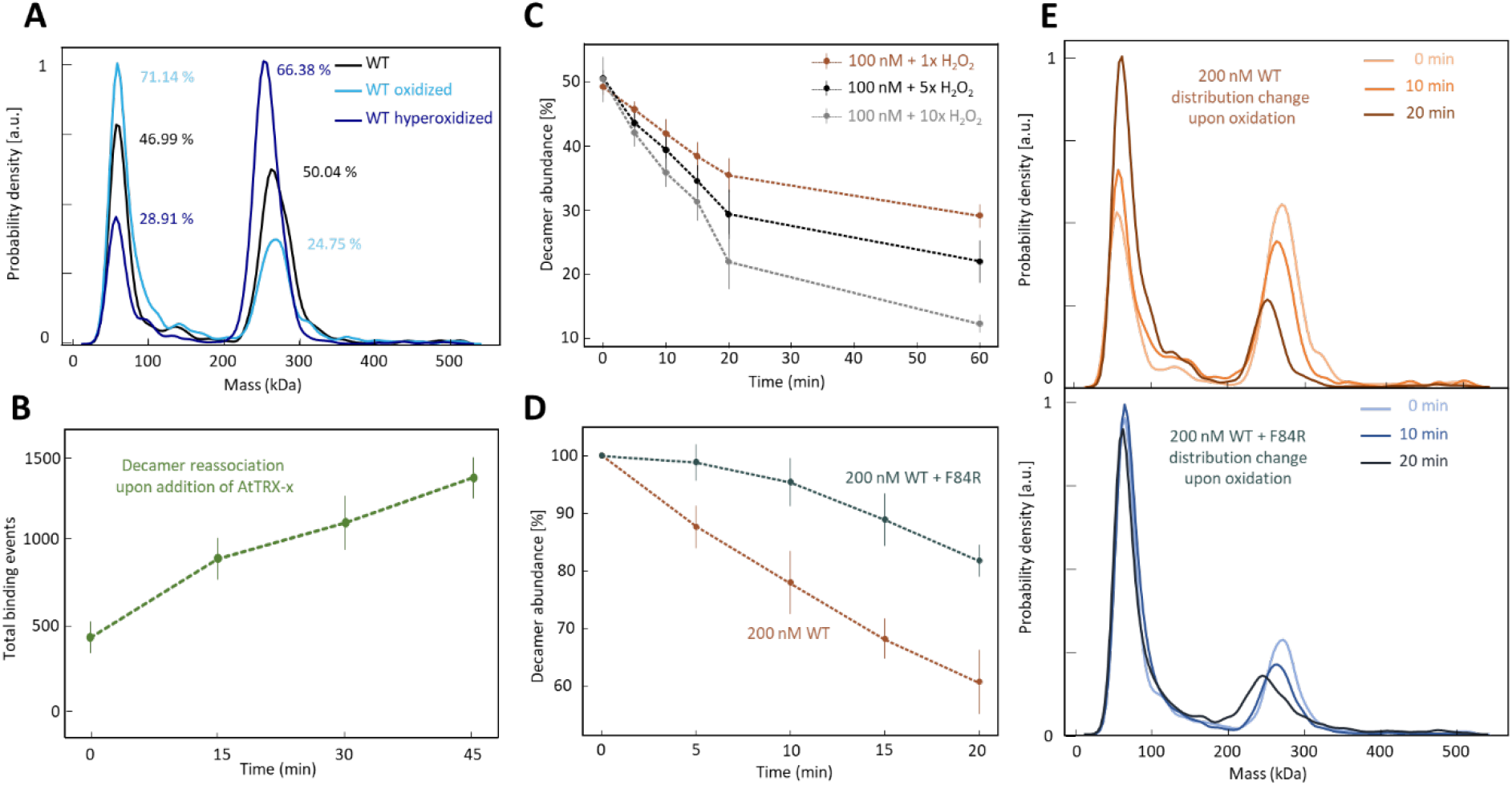
Thiol-oxidation induced changes in oligomerization state of At2-CysPRXA as visualized by single molecule mass photometry. (A) Distribution profiles of reduced (black line), oxidized (light blue) and hyperoxidized (dark blue) 2-CysPrxA at 100 nm concentration. Oxidation was achieved by incubating reduced 2-CysPRX with 1 μm H_2_O_2_ for 30 min and resulted in an increase of dimeric fraction. Hyperoxidation was induced by repetitive peroxidase turnovers of At2-CysPRXA by incubation with equal amounts of H_2_O_2_ and DTT (both 1 μm). Hyperoxidation promoted the formation of decamers relative to dimers. (B) Appearance of reduced At2-CysPRXA decamers after oxidation and regeneration by AtTRX-x in 5-fold excess as a function of time. The oxidized At2-CysPRX (t=0) was reduced using the cellular redox transmitter thioredoxin (TRX). The reduction increased the fraction of decamers 2-fold after 30 min. (C) Time-dependent oxidation of At2-CysPRXA by H_2_O_2_. H_2_O_2_ was added to 100 nm 2-CysPRX at equimolar (1x) or excess (5x or 10x) molar amounts. Oligomer distribution was monitored by mass photometry over 60 min. (D) Protection of At2-CysPRXA from oxidation by F84R. F84R only adopts the dimeric conformation with higher peroxidase activity. A mix of WT and F84R at a ratio 1:1 was exposed to 5-fold H_2_O_2_. The control assay contained twice the WT amount. Since F84R cannot decamerize, the detected decamer exclusively reveals the reduced WT form. Higher stability of WT oligomers in the presence of F84R indicates more efficient protection by the dimeric form. (E) Relative appearance of dimer and decamer in mass photometry distributions including At2-CysPRXA (upper diagram) and a mix of At2-CysPRXA and F84R. Statistics: All experiments were performed with n > 6 independent determinations on at least two different days. Relative and absolute decamer abundance was taken from single mass readings between 200 and 300 kDa. Data are mean values ±SD from *n>6* measurements on at least two independent days.

The oxidation-dependent dissociation of decamers was restored upon reduction with DTT but we did not observe this behavior for the hyperoxidized form that is insensitive to DTT (**Figure 3**). The hyperoxidized wildtype accumulated as decamer and decameric stacks similar to the C54D variant which mimics exactly this state (Figure 1D). Hyperoxidation is enhanced under abiotic stress like heat resulting in exclusive accumulation of decamers and hyperaggregates likely by a differential charge distribution on the protein surface. [5]

**Figure 3.**
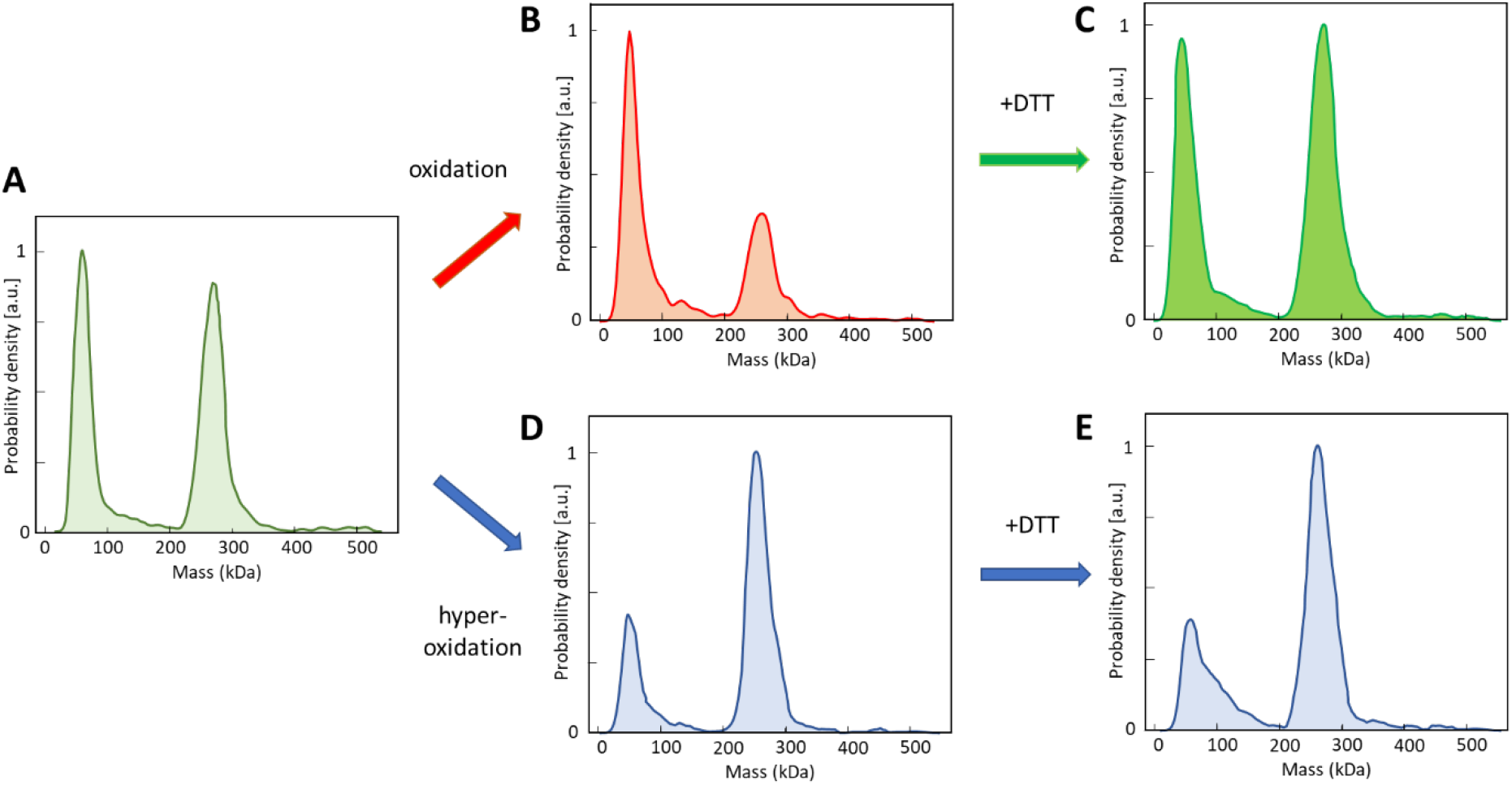
Distribution of At2-CysPRXA at 100 nm as affected by hyperoxidation and subsequent reduction. (A) Mass distribution of reduced 2-CysPRX from *Arabidopsis thaliana* at 100 nm concentration. (B) Mass distribution after oxidation on dimer:decamer distribution, (C) Effect of reduction by addition of 10-fold excess DTT. (D) Effect of hyperoxidation on dimer-to-decamer ratio by addition of 10-times excess DTT and H_2_O_2_ in equimolar amounts. (E) Inability of 10-fold excess DTT to restore reduced distribution shown in A. All protein samples were treated with redox additives in excess amounts 30 min before dilution to 100 nm. For readings each sample was kept on ice again for 20 min to ensure reproducibility and establishment of stable oligomer ratios. Statistics: Data are averaged values from *n>6* independent determinations performed at least at two different days.

Reductive regeneration of oxidized PRX in the cell is achieved by TRX or other electron donors. [24] Accordingly, adding excess AtTRX-x to oxidized At2-CysPRXA caused re-association of reduced decamers over time **(Figure 2B)**. All tested TRXs displayed a size of 40-60 kDa in mass photometry possibly due to a resolution limitation for smaller proteins like 10-12 kDa TRXs (**Supporting Figure 4**), and this readout for TRXs overlapped the 2-CysPRX dimer, impeding evaluation of 2-CysPRX dimers. Therefore, histograms were examined for higher oligomers specific for 2-CysPRX. Thereby, mass photometry allowed for monitoring redox transitions by counting single molecule populations even at very low concentrations in a unique way.

The results demonstrate our ability to rapidly determine the dependence of the oligomeric distribution of At2-CysPRXA on redox conditions. We therefore set out to quantify the time- and concentration-dependent oxidation in the presence of equimolar, 5x or 10x excess of H_2_O_2_ **(Figure 2C)**. In line with results from **Figure 2A** we found H_2_O_2_-dependent oxidation causing an increase in the dimer population at the expense of decameric species (Figure 2C). The initial dissociation of decamers was faster with elevated H_2_O_2_ (0-20 min) while a similar decrease in decamers was observed at later time points (20-60 min). Close inspection of the time-dependence of decamer decay revealed two-fold stimulated rates of disassembly at 10-fold increased H_2_O_2_ concentration, suggesting that oxidation is more rapid than disassembly of decamers into dimers.

Given that oligomers larger than dimers have been observed *in vivo,* we next explored the relative efficiency of dimers vs. decamers in H_2_O_2_ detoxification. [25] To do this, we used the F84R variant, which exclusively exists as a dimer (Figure 1D) and combined it at a 1:1 ratio with WT decamers, before adding a 5-fold excess of H_2_O_2_. The amount of WT decamer served as a convenient readout of excess H_2_O_2_. In the presence of F84R, we observed almost complete protection of the wildtype decamer from oxidative destabilization during the first 5 min, with the protective effect compared to a sample lacking F84R prevailing until the end of analysis **(Figure 2D)**. The corresponding histograms highlight the dissociation of decamers over time with a concomitant increase in dimers **(Figure 2E)**. This increase was altered for F84R addition due to the higher relative abundance of dimers compared to decamers and was considered in the calculations. The result demonstrates that the optimization of thiol peroxidase efficiency would have been possible during evolution by inhibiting decamer formation in case of At2-CysPRX.

### 2.4. Human 2-CysPRX isoforms differ in oligomerization dynamics

Our observation of oligomer-specific activity, and its dependence on redox conditions suggests that differences in oligomeric distributions may be connected to the function in general and also in the different human 2-CysPRX variants. Therefore, we studied three 2-CysPRXs encoded by the human genome. At 50 nm monomer concentration, HsPRX1 exhibited a similar oligomeric distribution as At2-CysPRXA and did not adopt the eicosameric conformation in the concentration range studied here (**Figure 4A**, **Supporting Figure 5**). For HsPRX2 and HsPRX3 we could not find evidence for decamers, reminiscent of F84R (Figure 3A).

**Figure 4.**
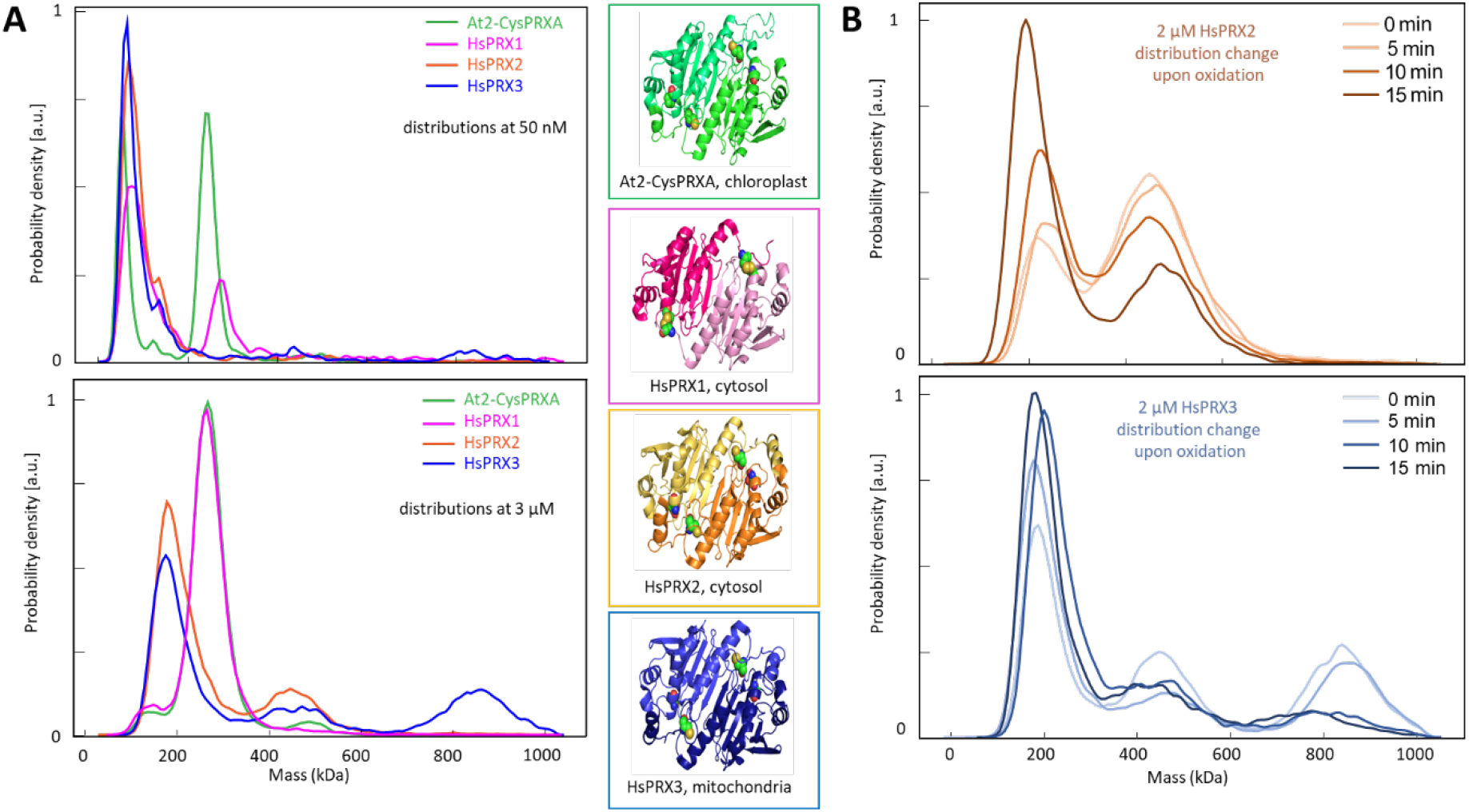
Diversification of oligomerization state of plant and human 2-CysPRX measured by single molecule mass photometry. (A) Oligomerization state of human peroxiredoxins HsPRX1 (cytosol, pink), HsPRX2 (cytosol, orange), and HsPRX3 (mitochondria, blue) and plant 2-CysPRXA (chloroplast, green). Distribution profiles are presented for low (50 nm, upper diagram) and high concentrations (3 μm, lower diagram). At2-CysPRXA and HsPRX1 shared the highest similarity in distribution whereas HsPRX2 and HsPRX3 remained in a more dimeric state at 50 nM. HsPRX2 and HsPRX3 adopted the conformation of higher order oligomers (eicosamer and eicosamer stacks (HsPRX3 only)) at 3 μm. (B) Effect of oxidation on the oligomization state of human 2-CysPRXs. HsPRX2 eicosamers dissociated upon oxidation with 5-fold excess H_2_O_2_ in a similar manner as plant 2-CysPRX. Oxidation of HsPRX3 caused a time-dependent dissociation of the higher order hyperaggregates. 3rd order oligomers dissociated faster than 2nd order oligomers. Statistics: Data are averaged values from n>6 independent determinations performed at least at two different days.

At 3 μm monomer concentration, HsPRX2/3 exhibited oligomer formation but compared to 50 nm only smaller multimers like tetra-, hexa- and octamers were formed (but not distinguishable) resulting in distributions tailing toward higher masses (Figure 3A, lower part). A high density of dimers may have the same effect of peak displacement. At higher concentration, we found evidence for decamer stacking (**Figure 4B**). For HsPRX3, a stack of four decamers dominated at 2 μm monomer concentration with eicosamers present as well. Similarly, we found signatures of eicosamers for HsPRX2, but little for HsPRX1. Taking monomer equivalents into stochiometric ratios, we see significant amounts of 2-CysPRX monomers stored in higher oligomeric structures for HsPrx2 and HsPrx3 (**Supporting Figure 6**). Oxidation by a 5-fold excess of H_2_O_2_ destabilized the decamer stacks resulting in an accumulation of smaller multimers, presumably dimers for both HsPRX2 and HsPRX3 (**Figure 4B)**. Oxidation of eicosamer and eicosamer stacks of HsPRX3 resulted in an unequal decay for both oligomers favoring the largest HMW stack for oxidation.

## 3. Discussion

Single molecule mass photometry allows assessing the conformational state and redox-dependent dynamics of 2-CysPRX in an unprecedented analytical depth. The results give new insight into the function of this abundant protein ubiquitously present in all cells. Previous concepts need to be revised based on the results of our study.

### 3.1. Dynamic oligomerization and the need for revising the critical transition concentration

Previous studies used ITC to explore the dissociation kinetics of At2-CysPRXA if injected at high concentration into buffer, e.g., 50 μm. The observed heat changes appeared in line with a model where the assumed decamer-dimer equilibrium had a dissociation constant (critical transition concentration) of 2.14 μm for At2-CysPRXA and 1.32 μm for human PRX1. The transition from significant heat release by dissociation to background signal, indicating ceased dissociation, occurred rapidly upon progressive injection. [15, 18] This interpretation needs revision since mass photometry revealed formation of decamers at low nanomolar concentration. An assumed low K_d_ (< 50 nm) indicates high abundance of oligomers under physiological conditions since the monomer concentration was estimated in planta with 100 μm in chloroplasts. [26] Such concentrations cannot be tested by mass photometry because single molecule events can no longer be resolved.

The second order oligomers derived from concentration-dependent analysis by mass photometry for decameric stacking is of the same order of magnitude as the critical transition concentration observed by ITC. Together with mass photometry analysis, this indicates that the ITC results likely describe the transition between the single decamer and decameric stacks, and not as previously assumed the dimer-decamer transition. However, the critical transition concentration as determined by ITC had a cooperativity or stoichiometric coefficient of about 130 which is not reflected in the transition between hyperaggregates and decamers. [15] Thus, a component of decamer/dimer dissociation may be involved as well, namely that upon disassembly of stable hyperaggregates, decamers dissociate into dimers according to the dissociation equilibrium observed above if injected into buffer.

### 3.2. Functional divergence of 2-CysPRX conformations

It was assumed that PRXs adopt five types of redox-dependent conformations namely reduced dimers, reduced decamers, oxidized dimers, hyperoxidized decamers and hyperoxidized stacks (Figure 1A). [2, 20] The results from mass photometry proves their existence but in addition demonstrates the presence of transition states such as tetramers and hexamers and also assemblies between decamers and subdecamers. Mass photometry gives no information on the stability of these transition states. However, the changing oligomerization behavior of the mutated 2-CysPRX variants shows the sensitivity of the oligomerization process to molecular features such as introduced charges in the catalytic center in C54D or at the dimer-dimer interface in F84R.

The dimer-only F84R variant has a 2.6-fold higher rate constant for H_2_O_2_-reduction than 2-CysPRX wildtype. [20] The improved performance of dimer-only 2-CysPRX could be demonstrated here since the presence of F84R protected wildtype 2-CysPRX from oxidation and decamer disruption (Figure 2D). This type of analysis was possible with mass photometry by counting the decamer fraction, since only wildtype can form decamers, whereas F84R cannot. This unique result underlines the functional significance of decamer formation as selective driving force in evolution since improved thiol peroxidase performance of the constitutive dimer could have easily be evolved by suppressing the ability for decamer formation for the plant 2-CysPRX. This may be different for HsPRX2 and HsPRX3 that were reported to display higher thiol peroxidase activity as decamers by stabilization of the active site loop-helix. [27] In plants, the differential peroxidase efficiency of dimers and decamers of At2-CysPRXA may be important to avoid its complete oxidation and to limit futile reduction-oxidation cycles in the cell.

The plant 2-CysPRX functions as TRX oxidase important to reverse reductive activation of photosynthetic enzymes. [8, 28] Essentially, this mechanism represents a futile cycle, since oxidized enzymes and TRXs need to be reduced again in order to keep the Calvin-Benson cycle active and was recently simulated by mathematical modeling. [29] Future models should include the presence of pools of differently efficient thiol peroxidases in order to estimate their significance for keeping the futile cycle in check. KM(H_2_O_2_)-values of bacterial and plant 2-CysPRX were reported with 1-2 μm. [26, 30] Thus, it is intriguing that 100 nm H_2_O_2_ was able to oxidize a major portion of At2-CysPRXA confirming the extremely high substrate affinity of 2-CysPRX.

The chaperone activity of the C54D variant is 4-fold higher than that of wildtype At2-CysPRXA. [20] Considering the transition of dimers to oligomers of reduced At2-CysPRXA at concentrations below 50 nm, the chaperone activity measurements in that study were dominated by decamers at 10 μm. Nevertheless, the C54D variant mimicking hyperoxidized decamers (Figure 1D) was more efficient in stabilizing citrate synthase at elevated temperature compared to the reduced decamers of wildtype.

Jang et al. proposed a chaperone function of the hyperoxidized HMW forms under conditions of oxidative stress. [5] Detailed studies in Arabidopsis, barley and potato revealed species- and isoform-specific variation in the susceptibility of 2-CysPRX to hyperoxidation. [31] The function of the hyperoxidized form as chaperone may be significant but does not explain the physiological advantage of the dimer-oligomer-multimer equilibria of the reduced form with far less chaperone activity. The reason for keeping the redox-dependent conformational dynamics is likely due to the specific interactions of 2-CysPRX with other proteins as revealed in a proteomic study. [32] As an example from this study, the C54D variant bound to β-carbonic anhydrase and inhibited its activity to 50% in an enzyme activity test. [32]

### 3.3. Functional conservation and specification in human 2-CysPRX in comparison to At2-CysPRX

The comparison of the oligomeric state of four 2-CysPRXs, namely At2-CysPRXA, HsPRX1, HsPRX2 and HsPRX3 revealed distinct distribution patterns of dimers, decamers, eicosamers and eicosamer stacks. The highest similarity was detected between At2-CysPRXA and HsPRX1 tentatively suggesting strong overlap in function as discussed above. A significant fraction of HsPRX1 adopts the decameric conformation at low concentration (Figure 4). It should be noted that these results were obtained with native recombinant protein expressed in *E. coli* where posttranslational modifications are essentially missing. Human peroxiredoxins undergo profound posttranslational modifications such as phosphorylation, glutathionylation and acetylation and it would be important to study the oligomerization state of posttranslationally modified variants by mass photometry as well. [33]

HsPRX3 regulates apoptosis in human cells. RNAi-mediated suppression of HsPRX3 accumulation sensitizes HeLa cells to staurosporine- or tumor necrosis factor alpha-induced apoptosis, while overexpression suppresses programmed cell death. [34] Furthermore, HsPRX3 was demonstrated to form stacks and tubes depending on pH and redox state. The authors termed this process self-chaperoning hinting at a particular function upon hyperoxidation. [35] Reduced HMW aggregates of HsPRX3, as shown with mass photometry here, may serve as storage pools to release dimers and decamers for efficient peroxide detoxification controlling apoptosis. The HsPRX3 content of a mitochondria-enriched protein fraction reached 1.9 μg monomer/mg protein in HeLa-cells. [34] Assuming a 25% protein solution, this corresponds to 23 μm HsPRX3 as monomer and 2.3 μm as decamer similar to the experimental condition used for Figure 3A.

Human 2-CysPRXs are implicated in many cell functions and diseases e.g., in cardiovascular and neurological disorder or carcinogenesis. [36, 37] Current understanding of PRX function suggests that PRX-dependent signaling processing involves formation of transient or stable complexes between target proteins, scaffold proteins and PRXs, like in the case of the regulation of the human transcription factors STAT3, which is oxidized by HsPRX2 and this interaction is stabilized by the presence of Annexin A2. [38] Mass photometry may prove important to understand the conformational requirements e.g., as dimers or decamers, of HsPRX2 to form such signaling domains.

Mass photometry is particularly sensitive at low concentrations and is able to reveal dynamic processes whereas crystallization uses high protein concentrations up to 1 mm and depends on conformationally fixed proteins. [35] The same applies to cryomicroscopy. Neither method can provide information on dynamic changes in conformation. Both methods are biased toward detecting large structures and stable assemblies. The application of artificial environments and non-physiological protein concentrations may be the reason why several structures for 2-CysPrx exist without functional annotation. [16]

## 4. Conclusions

Single molecule mass photometry quantifies individual molecular entities in a larger ensemble. The benefit is that (i) the method determines masses of each detected molecular entity and not averages, (ii) the obtained distribution peaks provide reliable masses if the mass exceeds 40 kDa, (iii) the speed of determination allows for time resolution of kinetic changes in fractions of minutes, (iv) low protein amounts are sufficient for this method and (v) labeling with fluorophores is unnecessary. Therefore, mass photometry is a powerful and unique method to visualize the conformational states of 2-CysPRX in a native state at low concentrations. This study on chloroplast 2-CysPRXA and three human 2-CysPRX revealed distinct features of these thiol peroxidases and provides insight into the redox-dependent transition between the oligomeric states. The previous assumption of a highly cooperative transition between dimer and decamer at the low micromolar concentration needs revision. Both At2-CysPRXA and HsPRX1 adopt the decameric state in the nanomolar concentration range. Transition states such as tetra- and hexamers and e.g., 14-mers appeared at low frequency as well. The results have functional implication since we can now assess the distribution of quaternary structures in solution at low concentrations without introducing labels. This type of analysis will grant access to studying the impact of binding partners or posttranslational modifications on 2-CysPRX aggregations and thus functional state.

## 5. Experimental Section

### Recombinant proteins

Wildtype and variants of At2-CysPRXA were generated by [20]. All recombinant proteins were expressed, purified and treated according to [18]. Dr. Thorsten Seidel (Bielefeld University) supplied the plasmids for expression of human HsPRX1, HsPRX2 and HsPRX3. [15] The recombinant proteins were reduced after purification by Ni-NTA-affinity chromatography with 10 mm DTT for 1 h. Samples were dialyzed 3 times for 4 h each in order to change the buffer and remove DTT and imidazole (35 mm HEPES, pH 8, 100 mm NaCl). Aliquots were snap-frozen in liquid nitrogen, stored at −80°C, and immediately thawed before each experiment. All proteins were prepared 20 min prior analysis.

### Mass Photometry

All proteins were reduced as described above and diluted with fresh and degassed buffer (35 mm HEPES, pH 8, 100 mm NaCl) if not noted otherwise. At least two different batches of purified protein were used for all studies. A standard protein solution was daily used for calibrating the contrast intensity to mass values. Movies were taken with frame averaging of five below 1 μm protein concentration and with frame averaging of two at 1 μm and higher. Movie files were analyzed using the DiscoverMP software (Refeyn, Oxford, UK). Raw contrast values were converted to molecular masses using the mass factor from calibration, and binding events counted with 5 kDa resolution. Binding events below 40 kDa were indistinguishable from background. Settings were adjusted according to the specific visualization requirements and with a background reading of buffer only. Statistics were done for multiple readings on a single day with absolute numbers and across several days with relative values only.

### *Dynamic light scattering* (DLS)

DLS was conducted in the Biochemistry Department of Oxford University. All protein samples were reduced and analyzed at 10 μm concentration in a NanoBrook Omni with the OmniSIZE DLS software.

## Acknowledgements

Plasmids for human PRX expression were provided by Dr. Thorsten Seidel. Wilena Telman generated and supplied further batches of selected PRX proteins.

Received: ((will be filled in by the editorial staff))

Revised: ((will be filled in by the editorial staff))

Published online: ((will be filled in by the editorial staff))

## Contributions

KJD who had the original idea and ML conceived the project with further input from MSK and PK. ML performed experimental work including protein purification and mass photometry analysis. MSK supervised data acquisition, processing and evaluation. ML, PK and KJD wrote the paper.

## Supporting Information

Supporting information is available online. **Supporting Figure 1:** Oligomer distribution of plant At2-CysPRXA at 100 nm concentration plotted as monomer equivalents. **Supporting Figure 2:** Reproducibility of oligomer distribution determined by mass photometry. **Supporting Figure 3:** Dynamic light scattering of At2-CysPRX using 10 μm wildtype, F84R, and C54D. **Supporting Figure 4** Mass photometry readings of plant thioredoxins. **Supporting Figure 5:** Relative abundance of oligomers of HsPRX1 as a function of concentration. **Supporting Figure 6:** Oligomer distribution of plant and human 2-CysPrx at 2 μm monomer concentration.

## Table of Contents

Mass photometry allows non-invasive and label-free visualization of protein size and oligomer distributions. Using this technique, the oligomerization dynamics of Arabidopsis and human 2-Cysteine peroxiredoxins, its sensitivity to redox components, and their distinct oligomer composition across different species was revealed. The results indicate a significant influence of oligomerization on protein-protein-interactions and functional diversification in the cellular environment.

### Graphical Abstract

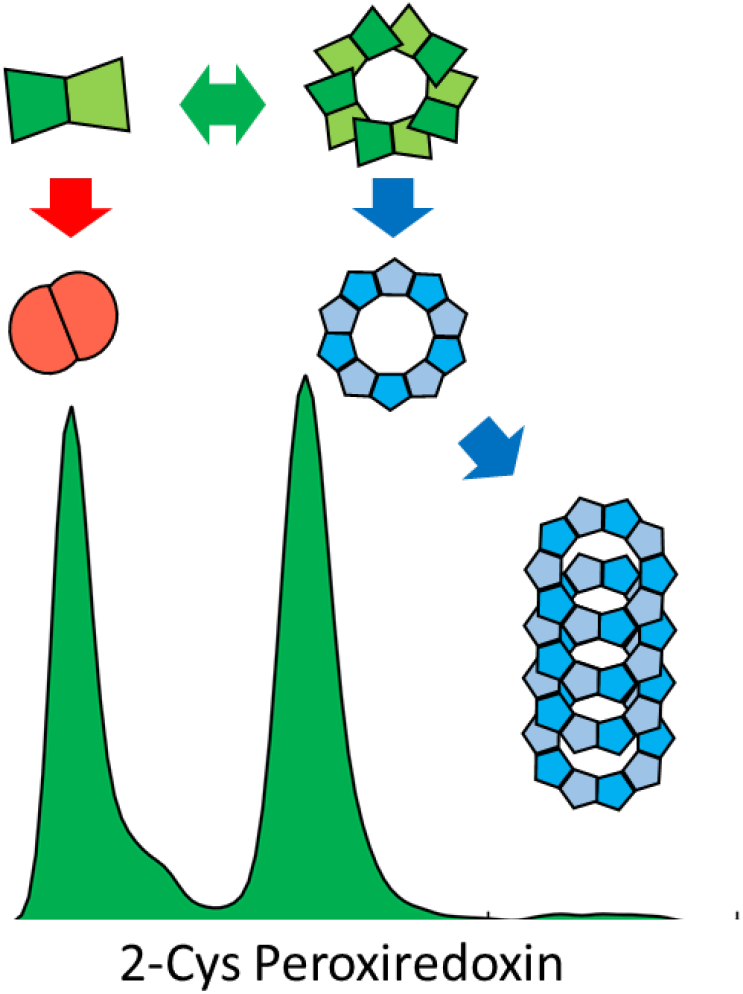

