## Supplemental materials Figures 1 - 6 for "Single Molecule Mass Photometry Reveals Dynamic Oligomerization of Plant and Human Peroxiredoxins for Functional Conservation and Diversification"

Supporting Fig. 1:
Liebthal et al.

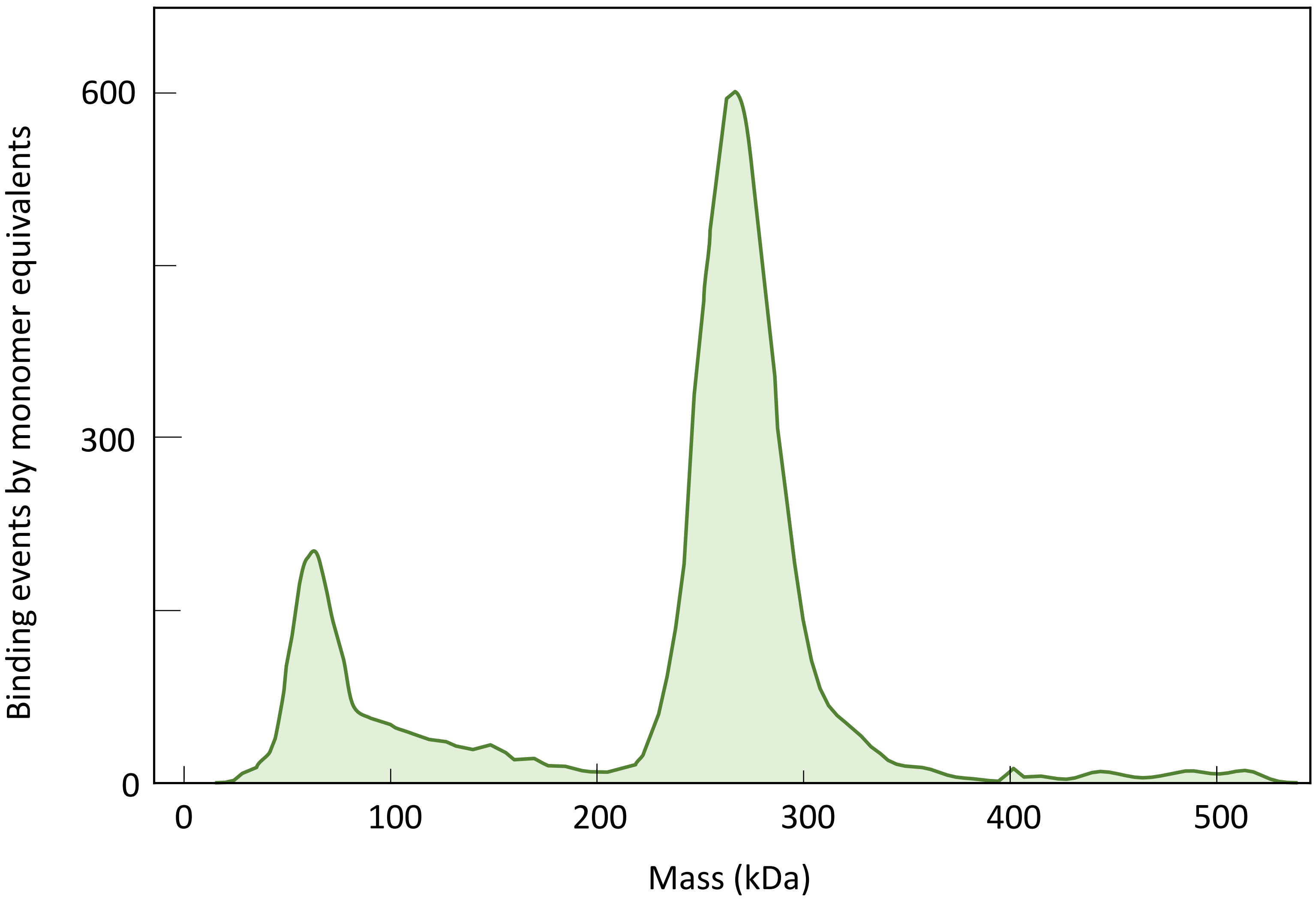


**Supporting Figure 1:** Oligomer distribution of plant At2-CysPRXA at 100 nM concentration plotted as monomer equivalents. Prior to analysis, the protein was reduced with DTT and diluted with degassed buffer (35 mM HEPES, pH 8, 100 mM NaCl). Recordings were taken after 20 min to exclude unequal dilution or oligomerization dynamics. Similar results were seen in n > 12 experiments. The plot was generated based on averaged values similar as presented in Supporting Figure 2.

Supporting Fig. 2:
Liebthal et al.

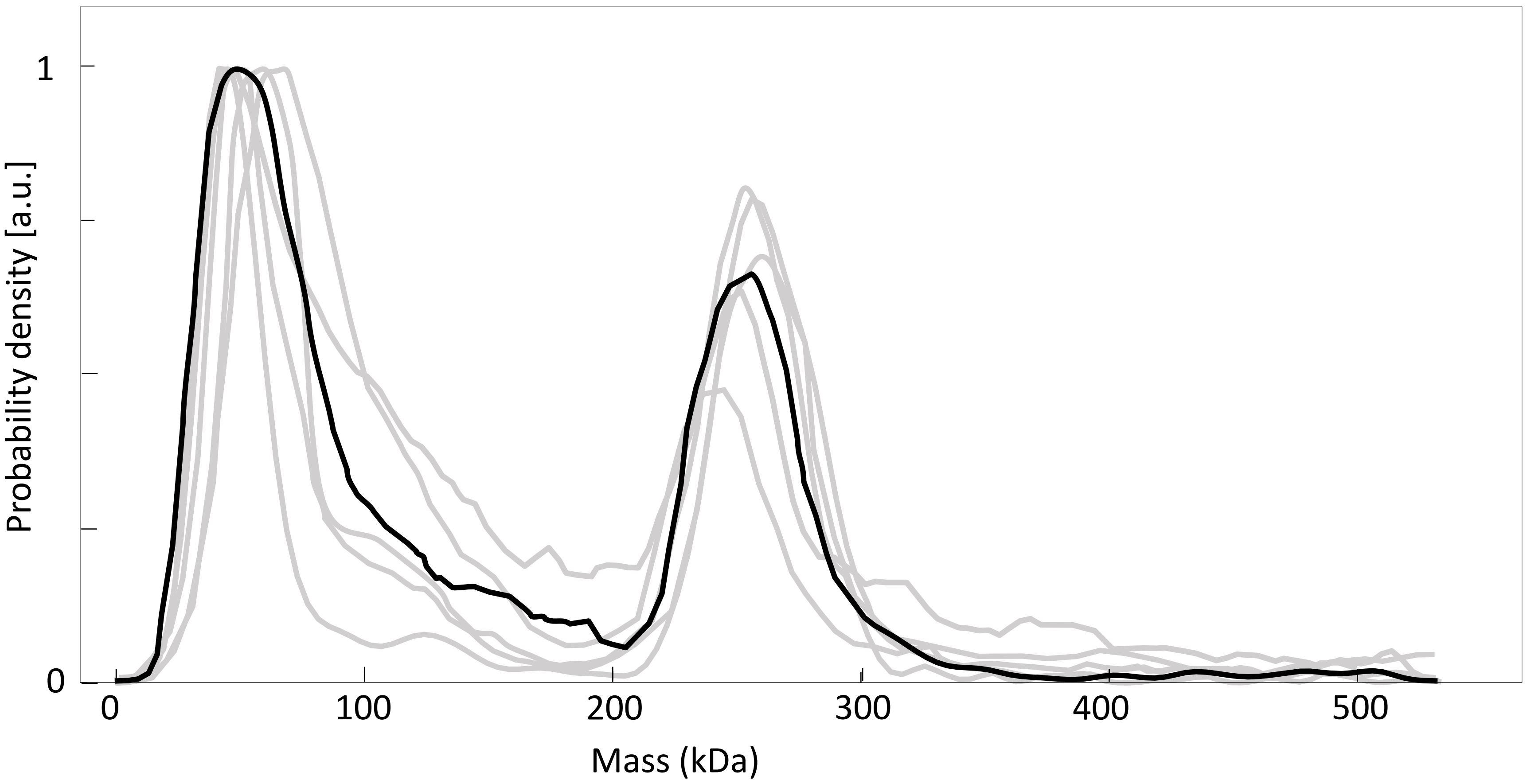


**Supporting Figure 2:** Reproducibility of oligomer distribution determined by mass photometry. The proteins were reduced with DTT and diluted with degassed buffer (35 mM HEPES, pH 8, 100 mM NaCl). Recordings were performed 20 min later to exclude unequal dilution or oligomerization dynamics. The overlay for 50 nM At2-CysPRXA consists of n = 5 independent readings (grey) and the averaged result (black) which is used in all figures.

Supporting Fig. 3:
Liebthal et al.

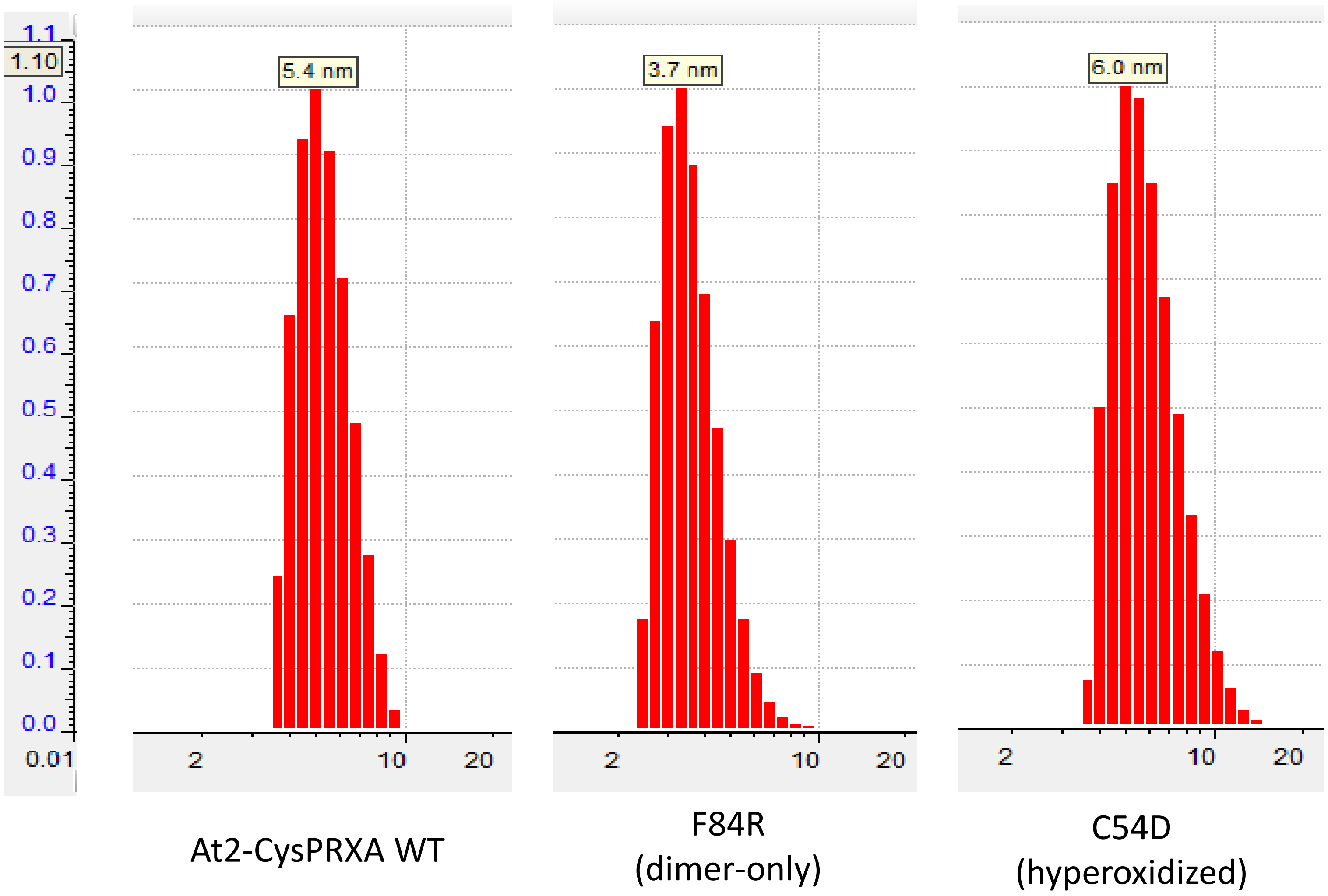


**Supporting Figure 3:** Dynamic light scattering of At2-CysPRX using 10 µM wildtype, F84R, and C54D. The samples were prepared as described for mass photometry and analyzed using a NanoBrook Omni with the OmniSIZE DLS software. Recordings were performed 20 min later to exclude unequal dilution or oligomerization dynamics. The dominant peak corresponding to the respective protein size (25 – 480 kDa) is depicted while other peaks describing impurities are excluded.

Supporting Fig. 4:
Liebthal et al.

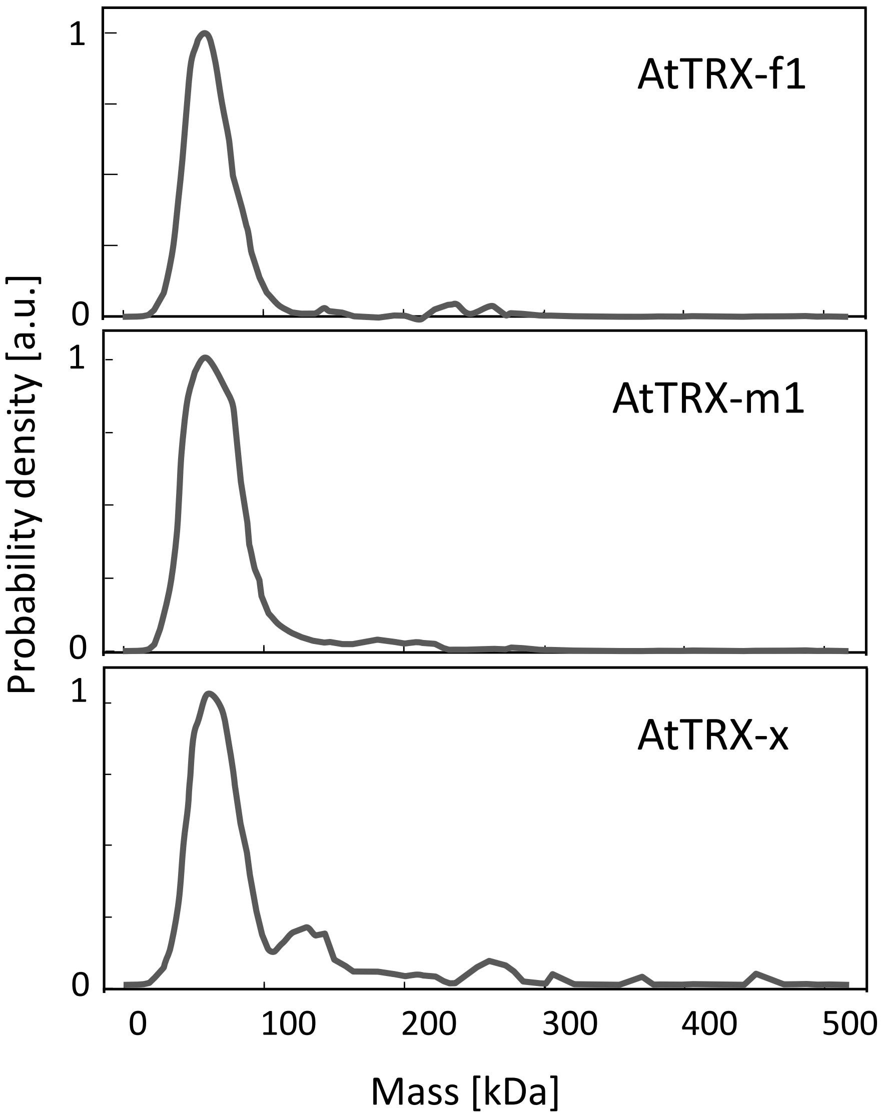


**Supporting Figure 4** Mass photometry readings of plant thioredoxins (TRX-f1/ TRX-m1/ TRX-x). The proteins were reduced by DTT, diluted with degassed buffer (35 mM HEPES, pH 8, 100 mM NaCl) and analyzed with mass photometry at 100 nM. Recordings were taken after 20 min. Data are means from n=3 independent readings.

Supporting Fig. 5:
Liebthal et al.

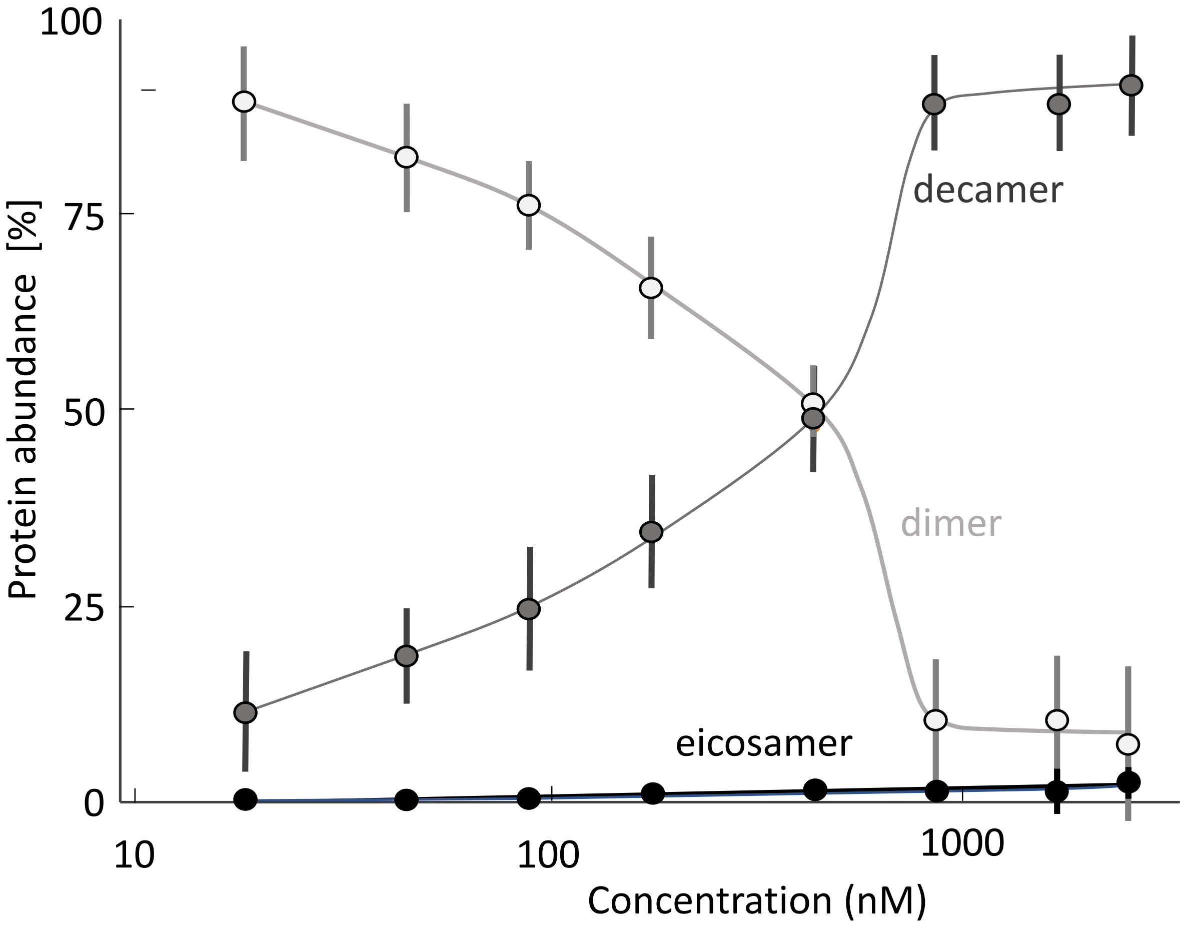


**Supporting Figure 5:** Relative abundance of oligomers of HsPRX1 as a function of concentration ranging from 20 nM to 3 µM. Proteins were reduced by DTT, diluted with degassed buffer (35 mM HEPES, pH 8, 100 mM NaCl) and analyzed with mass photometry. Recordings were taken after 20 min. The plots were generated based on averaged values similar as presented in Sup. Figure 2. To estimate to actual share in a distribution, the areas for dimers (0 to 100 kDa), decamers (200 to 300 kDa) and eisosamers (450 to 550 kDa) were integrated and compared to the total distribution area.

Supporting Fig. 6:
Liebthal et al.

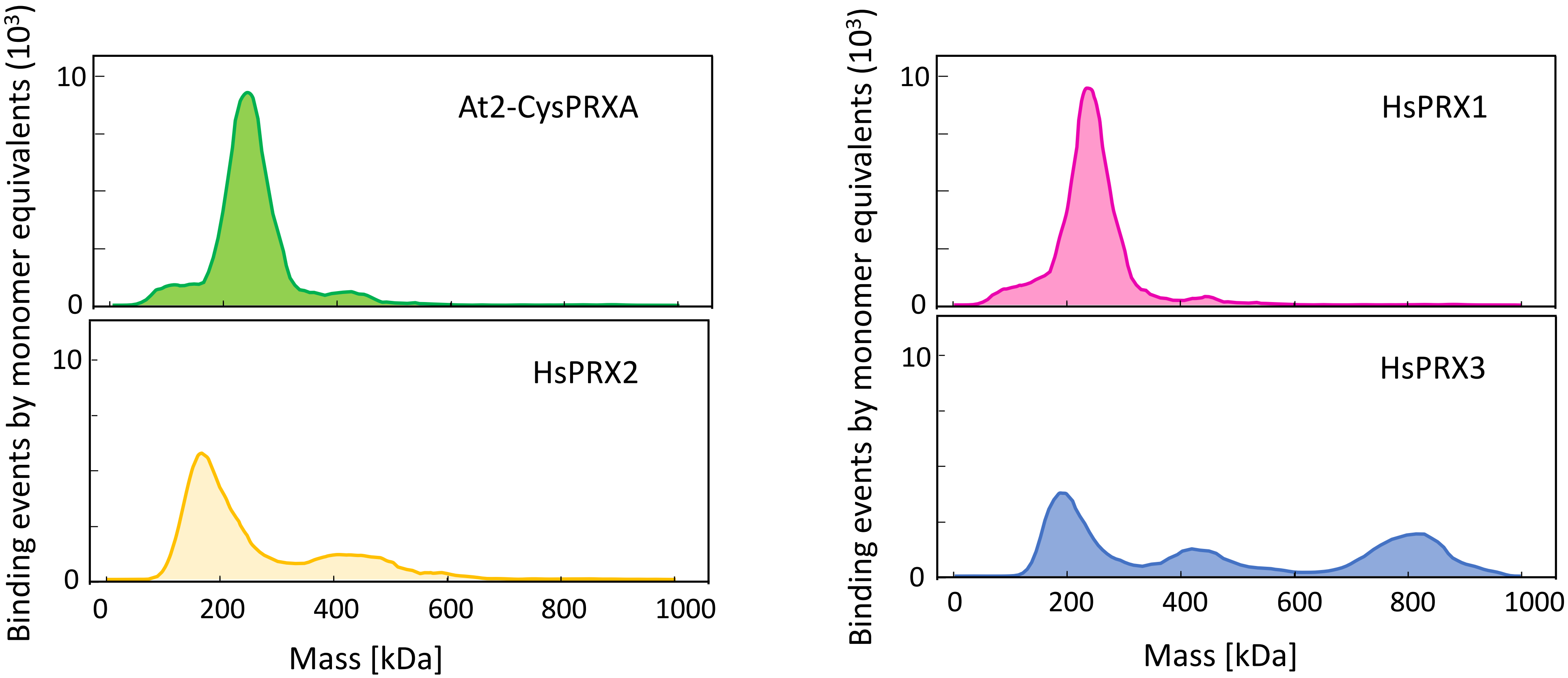


**Supporting Figure 6:** Oligomer distribution of plant and human 2-CysPrx at 2 µM monomer concentration. Recombinant proteins were analyzed by mass photometry at a concentration of 2 µM. Prior to analysis, the protein was reduced with DTT and diluted with degassed buffer (35 mM HEPES, pH 8, 100 mM NaCl). Measurements were done after 20 min. Similar results were seen in n > 8 experiments.
